# Biomass Production, Phosphorus Use Efficiency, and Fatty Acid Composition in *Phaeodactylum tricornutum* Cultivated in Wastewater from an Atlantic Salmon Recirculating Aquaculture System

**DOI:** 10.1101/2025.01.09.632295

**Authors:** Stian Borg-Stoveland, Vukasin Draganovic, Tove M. Gabrielsen, Kristian Spilling

## Abstract

Recirculating Aquaculture Systems (RAS) are becoming increasingly important as a sustainable method for fish production. However, further advancements are needed to enhance the sustainability of RAS, particularly in the management of waste-streams. Furthermore, the capital and operational expenditures of running a RAS is high, so solutions turning waste-streams into added value is a focus area for the industry moving forward. Wastewater is high in nitrogen, and phosphorus (P) is foreseen to be the limiting nutrient. Here we evaluated the growth dynamics, biomass production, nutrient uptake, phosphorus use efficiency (PUE), and fatty acid composition of the diatom *Phaeodactylum tricornutum* cultivated in aquaculture wastewater (AWW) from a land-based Atlantic salmon RAS. The diatom was cultivated in various treatments at pilot scale (10 L), including 100% AWW and AWW supplemented with silicon (Si) and P, and compared with a control treatment using Guillard’s F/2 medium. The highest dry weight biomass concentrations of ∼2.0 g L⁻¹ was achieved for the nutrient-supplemented AWW treatments, even outcompeting the control, demonstrating its viability as a growth medium when supplemented with Si and P. The control treatment achieved the highest specific growth rate of 0.422 d⁻¹. The “Si+P addition” treatment achieved the highest PUE, indicating that co-supplementation strategies can enhance phosphorus utilization. However, fatty acid yields, particularly eicosapentaenoic acid (EPA) and docosahexaenoic acid (DHA), were lower in AWW-based treatments, likely due to micronutrient deficiencies or the effect of possible contaminants. Despite this, *P. tricornutum* cultivated in AWW offers a sustainable and possible cost-effective solution for producing biomass and valuable fatty acids.

## 1. Introduction

Recirculating Aquaculture Systems (RAS) are designed to reduce diseases and improve sustainability of fish farming. In the last two decades, there has been a significant advancement in the technology of RAS, particularly in land-based operations [1]. The nutrients in RAS wastewater streams provide an opportunity for resource recovery, promoting sustainable practices in nutrient reuse and circularity. By recovering valuable nutrients such as nitrogen and phosphorus from the wastewater, it is possible to reduce dependency on synthetic growth medium and redirect these resources into e.g. industrial applications, creating closed-loop systems that enhance resource efficiency [2]. The integration of microalgae cultivation with RAS wastewater treatment has been proposed as a sustainable solution to address nutrient pollution while simultaneously producing valuable algal biomass [3].

The utilization of microalgae in wastewater treatment and biomass production has gained significant attention in recent years due to their potential in bioremediation and as a sustainable aqua-feed ingredient [4, 5, 6]. The diatom *Phaeodactylum tricornutum*, an established microalgal model species, has emerged as a promising candidate due to its high growth rates and the ability to thrive in diverse environmental conditions [7, 8, 9] as well as its high content of polyunsaturated fatty acids (PUFAs), particularly eicosapentaenoic acid (EPA) and docosahexaenoic acid (DHA), which are crucial for the health of both humans and fish. The addition of *P. tricornutum* in fish feeds increases the nutritional quality of the fish, promote better growth, immune function, and overall health [10, 11].

The specific composition of nutrients, including nitrogen (N), phosphorus (P), and silicon (Si), plays a critical role in microalgal growth and fatty acid biosynthesis [12]. For example, P is an essential element in ribosome biogenesis, and its demand increases proportionally with the growth rate of an organism [13]. P limitation impacts growth not only by setting the upper limit for the number of ribosomes, but also by increasing the costs of managing excess quantities of other elements (e.g. N and Si), which accumulate due to constrained ribosome genesis [14].

The concept of Resource Use Efficiency (RUE) has a long history in ecology and is traditionally expressed as the ratio of biomass produced to the amount of a resource consumed [15]. It is rooted in the study of how organisms optimize the use of limited resources (e.g. nutrients) for growth and reproduction [16], and the common nutrients in focus have mainly been nitrogen (NUE) and phosphorous (PUE) [17]. However, less attention was initially given to PUE [18], despite the critical role of P in cellular processes like energy transfer (ATP), nucleic acid synthesis, and membrane structure [19].

PUE has been predominantly studied in the context of agriculture, focusing on optimizing crop productivity in P-limited soils [20]. In contrast, the exploration of PUE in microalgal systems, particularly in relation to their nutrient uptake capabilities in wastewater treatment and biomass production, remains relatively unexplored. A high PUE indicates that microalgae can maximize biomass production while minimizing phosphorus input. This efficiency is particularly important in wastewater treatment application, where microalgae can utilize N and P from the wastewater to support growth and production of protein, carbohydrates and fatty acids [21]. Additionally, nutrient supplementation, particularly of Si and P, has been demonstrated to impact the fatty acid composition of microalgae, as Si is essential for the growth of diatoms, influencing their cell wall structure and biomass production [22], while P availability is crucial for lipid metabolism and fatty acid synthesis [23]. Although the cultivation of microalgae in wastewaters has been investigated the past decade, only a small portion of the literature focuses on use of RAS wastewater. In particular, only three studies investigating salmon RAS wastewater is available [24, 25, 26].

To address the growing need for sustainable aquaculture wastewater (AWW) management and enhanced microalgal biomass production, we assessed the effects of various nutrient treatments in AWW from a land-based Atlantic salmon RAS. We recently demonstrated that several microalgal species, including *P. tricornutum,* are able to grow in AWW from this salmon RAS facility [27]. In addition, targeted nutrient supplementation can significantly boost both biomass and fatty acids production in microalgae, making nutrient management an essential component of effective microalgal cultivation [28].

The aim of this study was to investigate how specific nutrient treatments influence growth, biomass production and fatty acid content at pilot scale cultivation of *P. tricornutum*, using AWW as base growth medium. To assess this, we (i): quantified PUE and examined its relationship with biomass production and fatty acid yields. This involved assessing how efficiently P contributed to biomass production and fatty acid synthesis. We also (ii): examined the impact of nutrient supplementation on the fatty acid composition and biomass production in *P. tricornutum*.

## 2. Materials and methods

### 2.1 Collection and treatment of RAS wastewater

Wastewater for microalgae cultivation was obtained from a pilot-scale land-based RAS operated by Ecofishcircle AS in Lyngdal, Norway [27]. The RAS facility maintained a water volume of 160 m³ and was used to rear Atlantic salmon (*Salmo salar*) at a stocking density of approximately 55 kg m^-3^, with the fish having an average weight of around 3.2 kg at the time of wastewater collection. The salmon were fed a commercial diet (BIOMAR Orbit EX 1000 70 mg) that consisted of 33.6% crude fat, 35.8% crude protein, and 3.3% crude fiber. On the day of wastewater retrieval, about 67 kg of feed was administered.

### 2.2 Experimental setup and test species

The growth dynamics in the various treatments were monitored over a 14-day period. There were six treatments: Control, 100% AWW, Batch-fed, Si-addition, P-addition, and a combined Si+P addition (Table 1). All treatments were conducted in triplicate. Growth was monitored using relative fluorescence, and nutrient dynamics were evaluated by measuring concentrations of NO_2_+NO_3_, PO_4_, NH_4_, and Si. A simple and cost-effective bag-setup for algal growth (Fig. 1) was constructed based on [29]. The polyethylene bags were with a 90 µm wall thickness (Art.no 25582, Cowab, Norway) and sealed using an impulse sealer (Art.no 25567, Cowab, Norway). Each bag was designed to hold 10L of liquid. The top part of each bag was folded over four times, then six evenly spaced cuts were made to allow a round wooden stick to be threaded through. This setup secured the bag to the rig. At the bottom of each bag, a sterile 1 ml pipette tip cut diagonally was inserted. The penetration point was sealed using parafilm and thread tape. A small piece of tubing was inserted into the pipette tip and secured using thread tape. It was then connected to a four-way valve that was used to regulate airflow and to retrieve samples during the experiment.

**Fig. 1.**
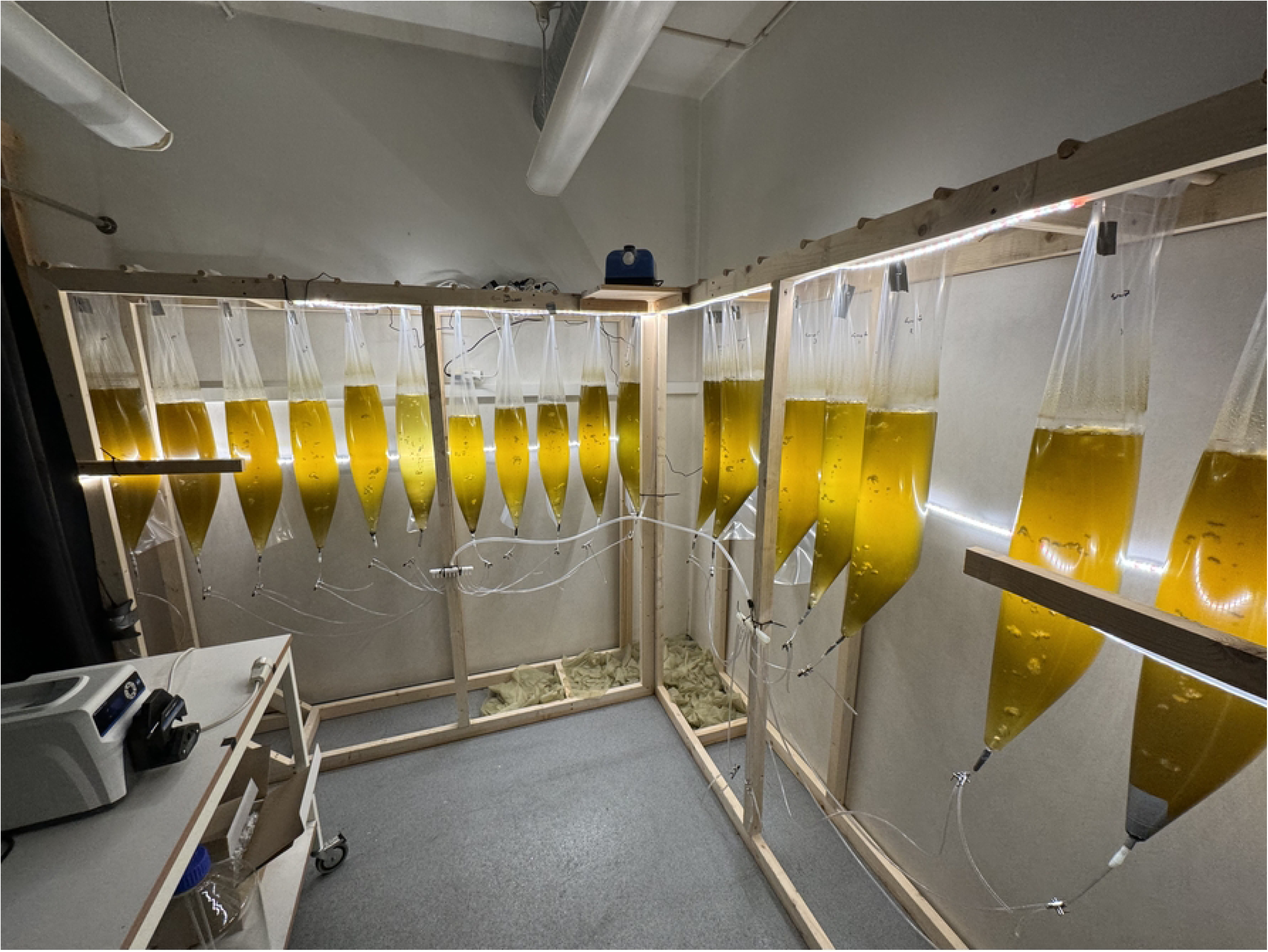
The setup on cultivation day seven. The cultivation bags supplied with air. The light strips were mounted behind the bags and on top of the rig. From the pump, the air was distributed to two ten-branch manifolds mounted on the two support beams. From there, the air was distributed to each bag via sterilized air hoses, at a rate of 2.5 L m^-1^

**Table 1:** Overview of the different treatments included in the experiment.

| Sample_ID | Treatment | Abbreviation | Sampling |
| --- | --- | --- | --- |
| G 1 (1,2,3) | Pure 1x F/2 + Si | Control | Nut + FA/OD |
| G 2 (1,2,3) | 100% AWW | 100% AWW | Nut + FA/OD |
| G 3 (1,2,3) | Pure AWW Batch-fed | Batch-fed | Nut + FA/OD |
| G 4 (1,2,3) | Pure AWW + Si | Si-addition | Nut + FA/OD |
| G 5 (1,2,3) | Pure AWW + P | P-addition | Nut + FA/OD |
| G 6 (1,2,3) | Pure AWW + Si & P | Si+P addition | Nut + FA/OD |

*Sample ID: numbers in the brackets (1,2,3) denotes replicates within one treatment. All treatments were conducted in 10 L volume except the Batch-fed treatment, where the initial volume was 5 L on day zero, and 1 L AWW was added every other day, until reaching 10 L on day 14. Nut = Nutrients, FA = Fluorescence, OD = Optical Density.*

For lighting, led-strips (Cotech, art.no 36-8517, China) providing a light intensity of ∼40 μmol m^-2^ s^-1^ (measured using a Li-Cor Li-1500, UK). Using a timer (Cotech 36-4600, type EMT757, China) light was supplied with a light/dark cycle of 16:8 h. The temperature was set to 15 °C. The bags were supplied with air from an air pump (AquaForte Air Pump HI-Flow V-60, AquaForte, Netherlands) at rate of 2.5 L m^-1^. From the pump, the air was distributed to two ten-branch manifolds, then via sterilized air hoses to a 4-way valve, both the valves and air hoses were sterilized in a 10% chlorine bath and then thoroughly rinsed in distilled water before use. The air supply was filtered through a 0.22 um filter (VWR, Avantor, item no. VWRI210-000126). One of the outlets on the 4-way valve was closed with a plug, one outlet directed air to the bag, and the last outlet was used to retrieve samples from the bags throughout the experiment. Filling of the liquid was done by making a small incision at the top of the bag, then using a peristaltic pump (Masterflex® L/S, model 77916-20, Avantor Fluid Handling, VWR) to transfer the liquid. Each bag was inoculated with 10% inoculum of the initial volume. In the treatments that required the addition of nutrients (groups 4, 5 and 6), this was done by pipetting through the same incision. For the treatments that required addition of Si and P we used Krystazil 40 (BIM Norway AS) and potassium dihydrogen phosphorus (KH_2_PO_4_) (Fybikon AS, Norway, item no. 16088), respectively. The nutrients were added on cultivation day eight, and the amounts added were calculated to correspond with the initial concentration of the given compound on day zero (1.04 mg L^-1^ for Si, and 0.94 mg L^-1^ for PO_4_). To prevent possible contamination and simultaneously securing gas exchange, the incisions were sealed with sterile cellulose stoppers (VWR, Avantor, item no. 391-0412).

*Phaeodactylum tricornutum* (strain CCMP630) was purchased from The Norwegian Culture Collection of Algae (NORCCA). Stock cultures were grown and kept in a climate chamber at 11 °C with a light/dark cycle of 16:8 h. Cultures were maintained in 50 mL sterile cell culture flasks (VWR, Avantor) using commercial 1x F/2 growth medium (Sigma-Aldrich, USA) with 1 ml L^-1^ added silicate (Krystazil 40, BIM Norway AS) [30]. Inoculation cultures were prepared in aerated (2.5 L m^-1^) 10 L flasks (Duran®, Avantor, VWR) and cultivated for two weeks before inoculation. Salinity was adjusted to match the AWW (22 ‰) using a hand-held refractometer (Fredriksen Scientific, Norway).

### 2.3 Sampling and harvesting of microalgae cultures

Sampling was conducted every other day throughout the experiment, starting on day 0. Nutrient samples were retrieved using 50 ml falcon tubes (Avantor, VWR) and stored dark at −20 °C until further analysis. The samples were analyzed for NO_2_+NO_3_, PO_4_, Si and NH_4_ using a nutrient analyzer (Skalar SAN++®Classic, Skalar Analytical B.V., Netherlands), with the following protocols: ISO 13395:1996 “Determination of nitrite nitrogen and nitrate nitrogen and the sum of both by flow analysis (CFA and FIA) and spectrometric detection” (NO₃ + NO₂), ISO 15681-2: “Determination of orthophosphorus and total phosphorus contents by flow analysis, Part 2: Method by continuous flow analysis (CFA)” (PO₄), ISO-16264 “Water quality - Determination of soluble silicates by flow analysis (FIA and CFA) and photometric detection” (Si), “Fluorometric determination of ammonia in sea and estuarine waters by direct segmented flow analysis” [31] (NH_4_). Throughout the experiments, algal growth was tracked by assessing fluorescence (FA) and optical density (OD680), following the protocols outlined by [32]. FA and OD680 measurements were conducted using a portable Pulse-Amplitude-Modulation (PAM) fluorometer (AquaPen AP 110/C, Photon System Instruments, Czech Republic). Before each measurement, the samples were dark-adapted for 30 minutes to ensure that all light reaction centers were open. Note that on day six, the AquaPen used to measure fluorescence reached its measuring threshold, so from day eight and onwards the samples were diluted 10x using distilled H_2_O. The pH was monitored using a Inolab 7110 (Xylem Water Solutions Norway AS, Norway).

Prior to harvest, calcium hydroxide (Ca(OH)_2_) was supplied as a coagulant agent to the microalgal cultures at a concentration of 1.25 g L^-1^. Within 30 minutes most of the biomass was concentrated at the bottom of the sleeves and could easily be collected by simply removing the air supply tube and let the biomass flow by gravitation. The algal biomass was collected into sterile 800 mL cell culture flasks (Avantor, VWR) and left standing for another 30 minutes to further concentrate the biomass by flocculation. The concentrated biomass was then transferred to sterile 150 ml cell culture flasks (Avantor, VWR) using a peristaltic pump, and subsequently shipped to the accredited laboratories of Eurofins Food & Feed (Møllebakken 40, 1538 Moss, Norway) for freeze drying and fatty acids analysis (Eurofins internal freeze-drying code MO03U-1, fatty acids quantification code MO051-1). Dry weight was determined by using the gravimetric method (Eurofins internal code LW1XT), as part of the process prior to the analysis of fatty acids content.

### 2.4 Data treatment

R-studio [33] was used to organize and visualize the data. The specific growth rates (SGR) were calculated from the change in fluorescence in a determined time interval corresponding to the exponential growth phase according to the formula:

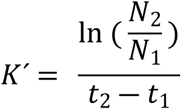

Where: *N_1_* and *N_2_* = fluorescence at time1 (*t_1_*) and time2 (*t_2_*), respectively [32]. Fluorescence data was log-normalized to better handle the skewness of the data.

## 3. Results

### 3.1 Growth and biomass production

Across all treatments, there was a general trend where algal growth increased towards day eight or ten, after which most treatments show either stabilization or a decline in growth by day 14 (Fig. 2). In the control treatment, algal growth increased steadily from day zero to eight, after which the biomass production plateaued. The specific growth rate (SGR) for the control treatment was the highest among all treatments (0.422 ± 0.005 d^-1^). In the 100% AWW treatment, algal growth was slower compared to the control at 0.285 ± 0.094 d^-1^. The Batch-fed, Si-addition and P-addition treatments exhibited similar growth patterns and SGRs (0.301 ± 0.029 d^-1^, 0.361 ± 0.015 d^-1^, 0.315 ± 0.002 d^-1^, respectively). In all three treatments, algal biomass increased steadily during the early phase, reaching a peak around day 10, followed by a plateau and a subsequent decline in fluorescence towards day 14. The combined Si+P treatment had a steady increase in fluorescence until day eight, with a peak observed on day 10, then followed by a sharp decline from day 10 to day 14. The SGR for the Si treatment was 0.267 ± 0.053 d^-1^, which was the lowest of all treatments.

**Fig. 2.**
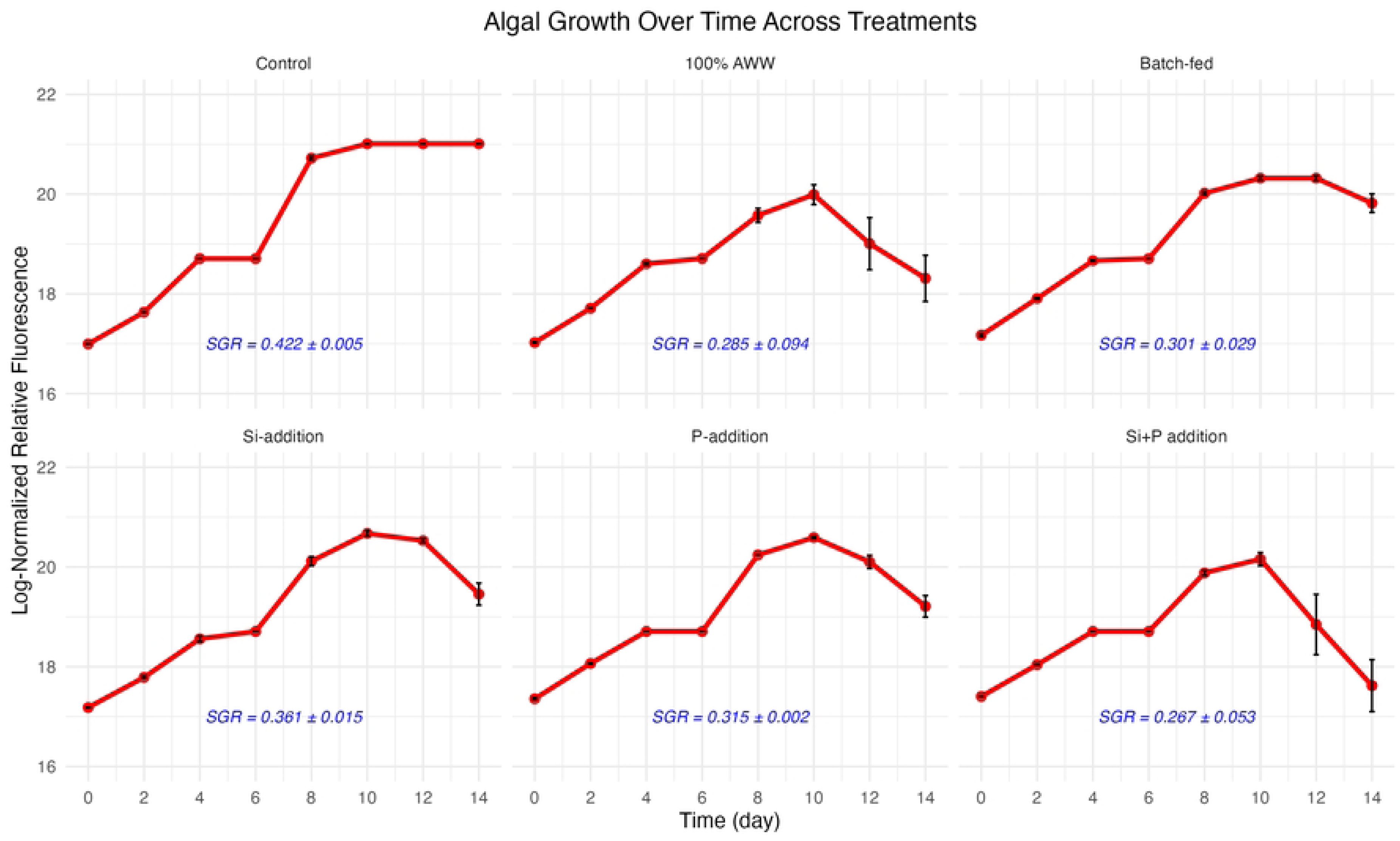
Growth dynamics (mean ± SE, n=3) of *P. tricornutum* over the 14-day cultivation period. Control is 100% Guillard’s 1x F/2 + added Si, while all other treatments were done with 100% AWW. Log-normalized relative fluorescence on the y-axis and cultivation time (days) is shown on the x-axis. For treatments “Si-addition”, “P-addition”, and “Si+P addition” nutrients were added on day eight.

The treatments Batch-fed, Si-addition and P-addition exhibited the highest biomass concentration after 14 days of cultivation, all three reaching around 2.0 g dw L^-1^ (Fig. 3). The Si+P addition treatment had a slightly lower biomass concentration, at 1.75 g dw L^-1^. The Control and 100% AWW treatments showed similar concentrations, but slightly lower between 1.5-1.7 g L^-1^.

**Fig. 3.**
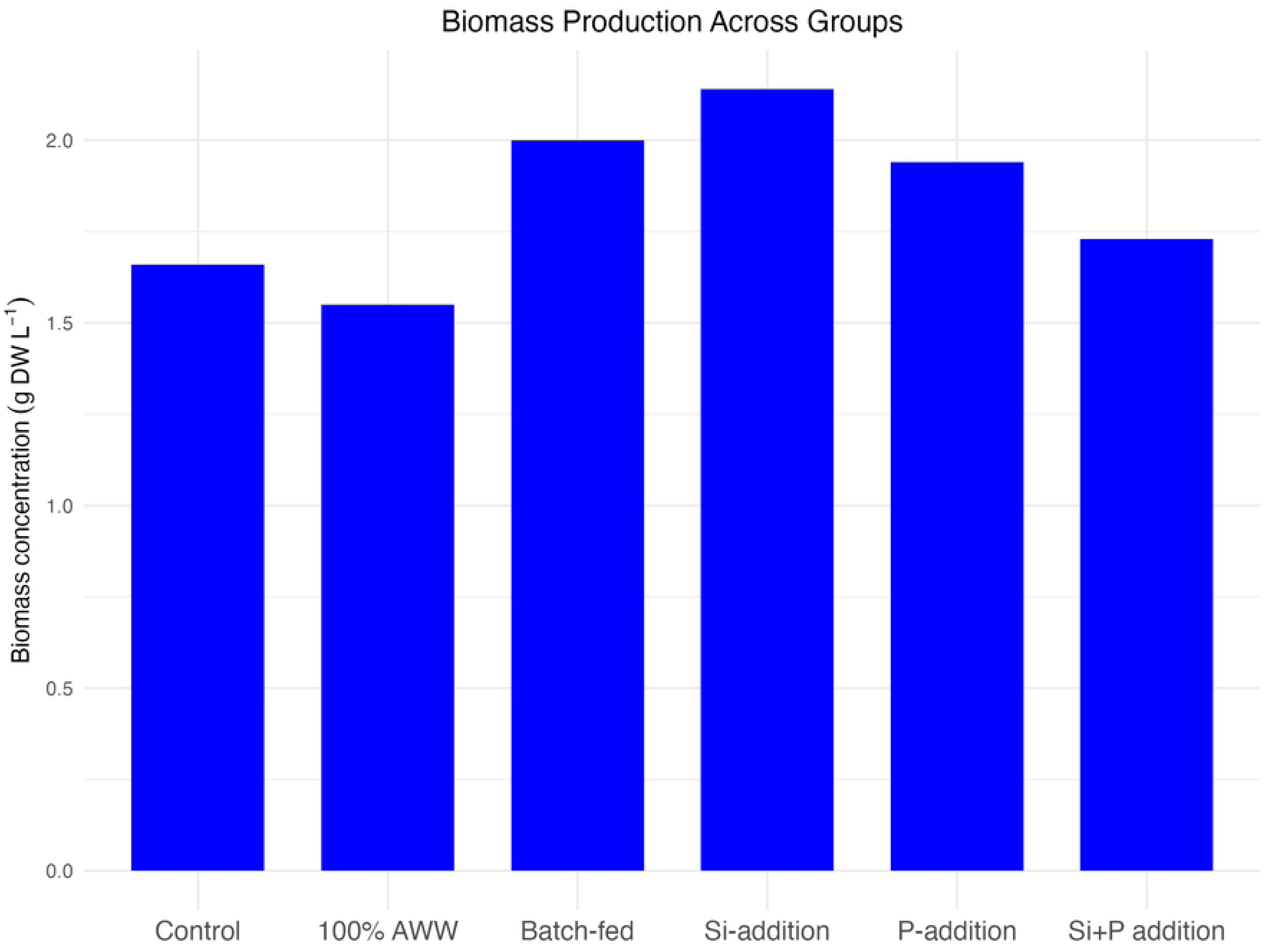
Biomass concentration across groups (g L^-1^) after 14 days of cultivation (n=1).

### 3.2 Nutrient uptake

In all treatments, including the control, nearly 100% of NO_2_+NO_3_, NH_4_ and PO_4_ were removed during the 14-day cultivation period (Table 2). The only exception being the P-addition treatment, where 79.5% of PO_4_ was removed. In the Batch-fed treatment, almost 95% of Si were removed, while the other treatments, excluding the control, showed an uptake between 35%-70%. However, the Control showed a modest uptake of 7.5%, however, note the initial concentration of 9.29 ± 0.62 mg L^-1^.

**Table 2:** Start (day 0), and end (day 14) concentrations for all compounds in the different treatments (mg L^-1^), mean ± SE).

| Compound (mg L<sup>-1</sup>) | Time | Control | 100% AWW | Batch-fed | Si-addition | P-addition | Si+P addition |
| --- | --- | --- | --- | --- | --- | --- | --- |
| NO<sub>2</sub>+NO<sub>3</sub> | Start | 12.31 ± 1.75(b) | 4.42 ± 0.21(b) | 4.42 ± 0.21(b) | 4.42 ± 0.21(b) | 4.42 ± 0.21(b) | 4.42 ± 0.21(b) |
|  | End | 0.00 ± 0.00 | 0.04 ± 0.05 | 0.03 ± 0.05 | 0.01 ± 0.02 | 0.15 ± 0.25 | 0.06 ± 0.0(a) |
|  | % Removal | 100% | 99.2% | 99.3% | 99.8% | 96.5% | 98.6% |
| NH<sub>4</sub> | Start | 0.00 ± 0.00(b) | 1.71 ± 0.38(b) | 1.71 ± 0.38(b) | 1.71 ± 0.38(b) | 1.71 ± 0.38(b) | 1.71 ± 0.38(b) |
|  | End | 0.00 ± 0.00 | 0.12 ± 0.16 | 0.03 ± 0.05 | 0.10 ± 0.17 | 0.00 ± 0.00(a) | 0.00 ± 0.00(a) |
|  | % Removal | N/A | 92.7% | 98.3% | 94.1% | 100% | 99.7% |
| PO<sub>4</sub> | Start | 1.08 ± 0.11 | 0.94 ± 0.04 | 0.89 ± 0.02 | 1.00 ± 0.04 | 0.91 ± 0.00 | 1.23 ± 0.03 |
|  | End | 0.00 ± 0.00 | 0.10 ± 0.17 | 0.00 ± 0.00 | 0.00 ± 0.00(a) | 0.19 ± 0.25(b) | 0.03 ± 0.00(a) |
|  | % Removal | 100% | 89.5% | 99.7% | 100% | 79.5 | 98.0% |
| Si | Start | 9.29 ± 0.62(b) | 1.04 ± 0.07(b) | 1.11 ± 0.08 | 1.10 ± 0.05 | 1.16 ± 0.05 | 1.12 ± 0.02 |
|  | End | 8.60 ± 0.94(b) | 0.48 ± 0.41 | 0.06 ± 0.10 | 0.33 ± 0.25 | 0.44 ± 0.47(b) | 0.72 ± 0.00(a) |
|  | % Removal | 7.5% | 53.7% | 94.6% | 69.8% | 62.2% | 35.4% |

Samples where n=1 is (a), n=2 is (b), n=3 is non-marked.

### 3.3 Phosphorus use efficiency (PUE)

In the treatments Control, 100% AWW, and Si+P addition, the PUE were around 1.5 g dw mg^-1^ P (Fig. 4). In the P-addition treatment, the PUE were almost double, reaching ∼2.7 g dw mg^-1^ P. Batch-fed and Si-addition displayed a PUE just above 2 g dw mg^-1^ P.

**Fig 4.**
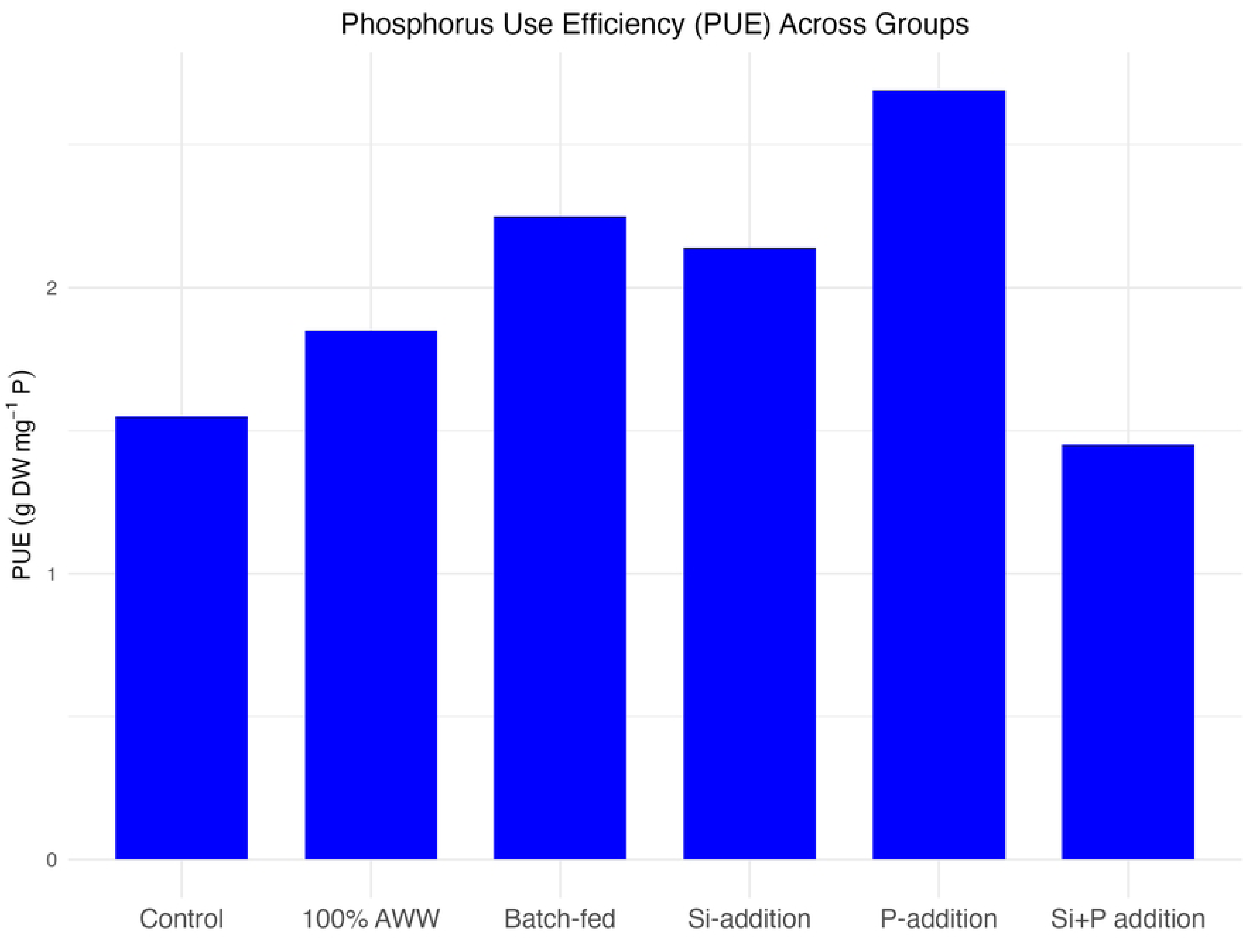
Phosphorus Use Efficiency (PUE). Phosphorus use efficiency across groups after 14 days of cultivation (g dw mg^-1^ P, n=1)

### 3.4 Fatty acids

The control treatment exhibited the highest fatty acids (FAs) yields overall. The total fatty acid yield was 16.58 mg g^-1^, with a substantial proportion coming from monosaturated fatty acids (MONOSATs) and saturated fatty acids (SATs), followed by polyunsaturated fatty acids (PUFAs) (Table 3). In the 100% AWW and Batch-fed treatments, the FAs yields overall were at approximately similar levels, with MONOSATs around 2 mg g^-1^ and SATs around 1.8 mg g^-1^. In the Si-addition and P-addition treatments, the yields dropped further, to around 1 mg g^-1^ for both MONOSATs and SATs, while n6-PUFAs and n3-PUFAs combined yielded approximately 0.25 mg g^-1^ and 0.2 mg g^-1^, respectively. In treatments Si-addition, P-addition and Si+P addition, the yields of n3-PUFAs, n6-PUFAs and PUFAs were around 0.2 mg g^-1^, 0.1 mg g^-1^, and 0.3 mg g^-1^, respectively. However, the yields of MONOSATs and SATs were approximately twice as high in Si-addition and P-addition compared to Si+P addition.

**Table 3:** Fatty acid yield (mg per gram dw) showing the major categories.

| Treatment | mg g<sup>-1</sup> dry weight |  |  |  |  |  |  | Total |
| --- | --- | --- | --- | --- | --- | --- | --- | --- |
|  | SATs | MONOSATs | PUFAs | n6-PUFAs | n3-PUFAs | EPA | DHA |  |
| Control | 3.86 | 4.77 | 3.37 | 0.47 | 2.15 | 1.82 | 0.14 | 16.58 |
| 100% AWW | 1.71 | 2.07 | 1.21 | 0.22 | 0.68 | 0.59 | 0.02 | 6.50 |
| Batch-fed | 1.82 | 1.88 | 0.69 | 0.13 | 0.39 | 0.33 | 0.02 | 5.26 |
| Si-addition | 1.17 | 1.11 | 0.29 | 0.06 | 0.17 | 0.14 | 0.01 | 2.95 |
| Pi-addition | 1.00 | 0.97 | 0.27 | 0.05 | 0.16 | 0.12 | 0.02 | 2.59 |
| Si+P addition | 0.49 | 0.49 | 0.28 | 0.05 | 0.16 | 0.11 | 0.02 | 1.60 |

*Total saturated (SATs), total monosaturated (MONOSATs), total polyunsaturated (PUFAs), n6-PUFAs, n3-PUFAs, eicosapentaenoic (EPA) and docosahexaenoic (DHA) (n=1)*

The control treatment also showed the highest yields of eicosapentaenoic acid (EPA) and docosahexaenoic acid (DHA) at 1.82 mg g^-1^ and 0.14 mg g^-1^, respectively. The DHA yield in all other treatments were quite modest between 0.01 and 0.02 mg g^-1^, and an EPA yield of 0.59 mg g^-1^ in 100% AWW, 0.33 mg g^-1^ in Batch-fed, and around 0.15 mg g^-1^ for Si-addition, P-addition and Si+P addition.

## 4. Discussion

The results of the present study provide evidence for the viability of using real aquaculture wastewater from a pilot scale land-based salmon farming as a growth medium for *P. tricornutum,* bringing AWW remediation one step closer to commercial use. Even though AWW alone can sustain growth, nutrient supplementation is necessary to optimize biomass production and PUE. The need for strategic nutrient addition is essential to fully leverage the potential of AWW as a sustainable resource in microalgae cultivation.

### 4.1 Growth dynamics and biomass production in AWW-based treatments

The specific growth rates (SGR) observed across treatments showed distinct differences between the control and the AWW-based treatments. The control treatment exhibited the highest SGR (0.422 d⁻¹), reflecting the optimized conditions of the F/2 medium, which provides balanced nutrients specifically designed to promote microalgal growth [30]. In contrast, the 100% AWW treatment exhibited a notably lower SGR (0.285 d⁻¹). This indicates that even though AWW contains essential nutrients for diatom growth, it may not provide optimal concentrations, or the balance required for *P. tricornutum* to achieve its full growth potential.

One possible inhibitory factor could be the NH₄ concentration, commonly found in wastewater, which can become toxic at elevated levels [34, 35]. However, when [35] tested NH₄ tolerance across six classes of microalgae, they found that concentrations below 100 µM were unlikely to inhibit microalgal growth. Given that the NH₄ concentration in the AWW used in this study was 0.27 mg L⁻¹ (∼19 µM), it is unlikely this level had any inhibitory effect on growth. In addition to NH₄ toxicity, certain heavy metals and excess trace nutrients can cause toxicity or interfere with nutrient uptake, leading to an imbalance in essential growth factors. For instance, even though iron is critical for diatom photosynthesis and nitrogen assimilation, excess levels can disrupt cellular redox balance and damage key metabolic pathways [36]. Heavy metals analysis was not conducted in this study. However, this limitation could have influenced the observed results, as unaccounted heavy metal toxicity might have contributed to growth imbalances [37]. Future studies should include heavy metal analysis to better understand their potential impact on nutrient uptake and metabolic functions.

Two of the nutrient-supplemented AWW treatments (Si-addition and P-addition) showed improved growth dynamics compared to untreated AWW, with SGR values of 0.361 d⁻¹ and 0.315 d⁻¹, respectively. These results suggest that the supplementation of critical nutrients like Si and P can partially compensate for the nutrient limitations in AWW, promoting better growth. The improved performance in these treatments highlights the role of Si in enhancing diatom growth, consistent with studies by [38], which emphasize Si’s importance for diatom cell wall formation and division. However, it has been shown that *P. tricornutum* is not dependent on Si for growth in a controlled environment [39]. Conversely, [40] showed that Si played an important role in the growth of this species under varying conditions. More recently, this was supported by [41] who found that high-silicate medium increased the production of biomass, and that such a strategy could be beneficial for large-scale cultivation. The fact that Si-supplemented AWW supported a higher growth rate compared to untreated AWW here, further underscores that Si plays a role in the growth of *P. tricornutum* in sub-optimal cultivation conditions. Our results also demonstrate the potential of AWW as a sustainable growth-medium alternative for large scale operations, and that targeted supplementation strategies could be beneficial.

While control treatment had the highest specific growth rate, the AWW based treatments, Batch-fed, Si-addition, and P-addition achieved comparable biomass yields (around 2.0 g dw L⁻¹). This is a crucial finding, as it demonstrates that with appropriate nutrient supplementation, salmon AWW can be a viable growth medium for *P. tricornutum*, achieving biomass yields surpassing those obtained with synthetic media. In commercial production, the biomass yield is a key economic parameter when evaluating profitability at scale. This finding aligns with other studies showing that microalgae can thrive in wastewater environments, especially when supplemented with key nutrients. For example, [42] demonstrated that wastewater can provide the necessary nutrients to sustain algal growth, but nutrient supplementation, particularly with N, P, and Si, can significantly enhance production.

The biomass production achieved in this study further supports that AWW, when combined with supplemental nutrients, can be a cost-effective and sustainable medium for large-scale microalgal cultivation [43]. The results from the 100% AWW treatment (∼ 1.5-1.7 g dw L⁻¹) indicate that while AWW alone is capable of supporting growth, the addition of specific nutrients such as P and Si is necessary to enhance biomass production.

### 4.2 Phosphorus use efficiency (PUE)

The phosphorus use efficiency (PUE) is defined as the ratio of the biomass produced to the bioavailable phosphorus consumed. This means that PUE is not directly equivalent to the biomass yield, as it specifically considers the phosphorus that is consumed, rather than the total phosphorus input. PUE is linked to the stoichiometric P ratios of the biomass, and increasing PUE is positively correlated to the DW : P or C : P ratios of the biomass, but not necessarily 1:1 as there for example might be uptake of inorganic P that is released again e.g. as dissolved organic phosphorus (DOP). In a system where phosphorus is the limiting nutrient, the biomass produced will be positively correlated to the PUE. However, if P is not the limiting nutrient, the relationship between inorganic P drawdown and biomass would be more complex.

In our results, the highest PUE was observed in the treatment with added phosphorus, which was the only treatment with measurable phosphorus concentrations at the end of the growth period. Interestingly, this suggests a relatively lower DW : P ratio in the biomass under phosphorus-limited growth, which appears counterintuitive. While we only had one replicate of PUE per treatment and cannot draw definitive conclusions, one possible explanation could be more active phosphorus uptake and storage during periods when phosphorus availability was declining.

### 4.3 Fatty acid yield and composition across treatments

Our results show that the control treatment yielded the highest overall fatty acid content, including the highest levels of EPA (1.7 mg g⁻¹). In contrast, all other treatments using 100% AWW with various nutrient supplementations, produced notably lower EPA and DHA yields. These findings correspond with the findings by [12], which reports that nutrient-rich synthetic media typically support higher fatty acid yields in *P. tricornutum*, especially for EPA and DHA.

The reduced yields of EPA and DHA in the AWW-based treatments indicate that while AWW can support the growth of *P. tricornutum*, the nutrient profile of AWW may not be optimized for fatty acid biosynthesis. In particular, the lower availability of micronutrients in AWW (e.g., sulfur, iron, calcium, zink and copper) could be limiting factors in the production of PUFAs [44]. Another potential factor is the presence of organic and inorganic contaminants in AWW that may interfere with metabolic processes. [45] investigated the use of microalgae for the treatment of municipal wastewater, and while their focus was not exclusively on AWW, they demonstrated that contaminants such as heavy metals, which are often present in AWW, could affect the metabolic processes, potentially inhibiting lipid biosynthesis. Furthermore, [46] found that contaminants in the wastewater from shrimp farms altered the fatty acid composition, resulting in a different lipid profile compared to synthetic media.

However, despite the lower fatty acid yields in the AWW-based treatments, the ability of *P. tricornutum* to produce EPA and DHA under these conditions suggests that AWW can serve as a cost-effective and sustainable medium for producing these high-value compounds. The challenge, however, lies in optimizing the nutrient profile of AWW through targeted supplementation to boost fatty acid synthesis while simultaneously maintaining effective growth.

### 4.4 Implication for large-scale cultivation using AWW as a growth medium

Using AWW as a growth medium reduces the need for costly commercial media, which is particularly valuable in large-scale operations where nutrient input constitutes a major economic expense [43]. The PUE and biomass production rates observed in the Si+P supplemented AWW treatment, indicate that AWW, with targeted nutrient supplementation, can support substantial growth. Such a setup could yield biomass levels similar to those achieved in synthetic media, but at a significantly reduced nutrient input cost.

The integration of *P. tricornutum* cultivation into RAS facilities would be feasible within existing aquaculture systems, allowing nutrient-rich wastewater to be directly repurposed, reducing the need for effluent treatment. This could streamline regulatory compliance by lowering N and P loads in discharged water. For large-scale operations, the approach offers a direct path to sustainability and economic efficiency by using wastewater as a resource rather than treating it as a disposal issue.

## 5. Conclusions

This study demonstrates that *P. tricornutum* can be successfully cultivated in AWW, particularly when supplemented with key nutrients like Si and P. While AWW alone supported high biomass production, nutrient supplementation improved both growth dynamics and PUE, achieving biomass concentrations outcompeting F/2 media. The enhanced PUE observed in nutrient-supplemented treatments underscores the importance of balanced nutrient availability for maximizing growth and efficiency in microalgal cultivation. However, fatty acid yields, particularly EPA and DHA, were lower in AWW-based treatments compared to the control, likely due to micronutrient deficiencies and potential contaminants in AWW. Despite these challenges, AWW remains a promising and cost-effective medium for sustainable large-scale microalgal cultivation, with opportunities for further optimization to enhance both biomass production and fatty acid yields.

## 6. Acknowledgments

The authors wish to thank Beate Marlene Funk-Stray for analyzing the nutrient data. Thanks also to Kristoffer Monrad and Andrea Noche Ferreira for assisting in retrieval of seawater and wastewater. We also want to thank Pontus Felix Eriksson for assisting with the experimental setup.

## Notes

### Competing Interest Statement

The authors have declared no competing interest.

## References

1. Espinal CA, Matulić D. Recirculating aquaculture technologies. In: Goddek S, Joyce A, Kotzen B, Burnell GM, editors. Aquaponics Food Production Systems: Combined Aquaculture and Hydroponic Production Technologies for the Future. Cham: Springer International Publishing; 2019. p. 35–76.

2. Nagarajan D, Lee D-J, Chen C-Y, Chang J-S. Resource recovery from wastewaters using microalgae-based approaches: A circular bioeconomy perspective. Bioresource Technology. 2020;302:122817.

3. Ansari FA, Guldhe A, Gupta SK, Rawat I, Bux F. Improving the feasibility of aquaculture feed by using microalgae. Environmental Science and Pollution Research. 2021:1–24.

4. Gonçalves AL, Pires JC, Simões M. A review on the use of microalgal consortia for wastewater treatment. Algal Research. 2017;24:403–15.

5. Carneiro WF, Castro TFD, Orlando TM, Meurer F, de Jesus Paula DA, Virote BdCR, et al. Replacing fish meal by Chlorella sp. meal: Effects on zebrafish growth, reproductive performance, biochemical parameters and digestive enzymes. Aquaculture. 2020;528:735612.

6. Rahman KM. Food and high value products from microalgae: Market opportunities and challenges. In: Microalgae biotechnology for food, health and high value products. 2020:3–27.

7. Butler T, Kapoore RV, Vaidyanathan S. Phaeodactylum tricornutum: a diatom cell factory. Trends in biotechnology. 2020;38(6):606–22.

8. Jiang J, Huang J, Zhang H, Zhang Z, Du Y, Cheng Z, et al. Potential integration of wastewater treatment and natural pigment production by Phaeodactylum tricornutum: Microalgal growth, nutrient removal, and fucoxanthin accumulation. Journal of Applied Phycology. 2022:1–12.

9. Russo MT, Rogato A, Jaubert M, Karas BJ, Falciatore A. Phaeodactylum tricornutum: An established model species for diatom molecular research and an emerging chassis for algal synthetic biology. Journal of Phycology. 2023;59(6):1114–22.

10. Cui Y, Thomas-Hall SR, Schenk PM. Phaeodactylum tricornutum microalgae as a rich source of omega-3 oil: Progress in lipid induction techniques towards industry adoption. Food chemistry. 2019;297:124937.

11. Uzlaşır T, Selli S, Kelebek H. Spirulina platensis and Phaeodactylum tricornutum as sustainable sources of bioactive compounds: Health implications and applications in the food industry. Future Postharvest and Food. 2024;1(1):34–46.

12. Yaakob MA, Mohamed RMSR, Al-Gheethi A, Aswathnarayana Gokare R, Ambati RR. Influence of nitrogen and phosphorus on microalgal growth, biomass, lipid, and fatty acid production: an overview. Cells. 2021;10(2):393.

13. Elser J, Acharya K, Kyle M, Cotner J, Makino W, Markow T, et al. Growth rate–stoichiometry couplings in diverse biota. Wiley Online Library; 2003.

14. Jeyasingh PD, Goos JM, Lind PR, Roy Chowdhury P, Sherman RE. Phosphorus supply shifts the quotas of multiple elements in algae and Daphnia: ionomic basis of stoichiometric constraints. Ecology Letters. 2020;23(7):1064–72.

15. Hodapp D, Hillebrand H, Striebel M. “Unifying” the concept of resource use efficiency in ecology. Frontiers in ecology and evolution. 2019;6:233.

16. Vitousek PM, Howarth RW. Nitrogen limitation on land and in the sea: how can it occur? Biogeochemistry. 1991;13:87–115.

17. Sheriff DW, Margolis HA, Kaufmann MR, Reich PB. 5 - Resource Use Efficiency. In: Smith WK, Hinckley TM, editors. Resource Physiology of Conifers. San Diego: Academic Press; 1995. p. 143–78.

18. Veneklaas EJ, Lambers H, Bragg JG, Finnegan PM, Lovelock CE, Plaxton WC, et al. Opportunities for improving phosphorus-use efficiency in crop plants. The New phytologist. 2012;195 2:306–20.

19. Canfield DE, Kristensen E, Thamdrup B. The phosphorus cycle. Advances in marine biology. 48: Elsevier; 2005. p. 419–40.

20. Richardson AE, Lynch JP, Ryan PR, Delhaize E, Smith FA, Smith SE, et al. Plant and microbial strategies to improve the phosphorus efficiency of agriculture. Plant and Soil. 2011;349(1):121–56.

21. Su Y. Revisiting carbon, nitrogen, and phosphorus metabolisms in microalgae for wastewater treatment. Science of the total environment. 2021;762:144590.

22. Yang M, Xu X-Y, Hu H-W, Zhang W-D, Ma J-Y, Lei H-P, et al. Combined application of nitrogen, phosphorus, iron, and silicon improves growth and fatty acid composition in marine epiphytic diatoms. Frontiers in Marine Science. 2023;10:1292713.

23. Zulu NN, Zienkiewicz K, Vollheyde K, Feussner I. Current trends to comprehend lipid metabolism in diatoms. Progress in lipid research. 2018;70:1–16.

24. Böpple H, Kymmell NLE, Slegers PM, Breuhaus P, Kleinegris DMM. Water treatment of recirculating aquaculture system (RAS) effluent water through microalgal biofilms. Algal Research. 2024;84:103798.

25. Böpple H, Slegers PM, Breuhaus P, Kleinegris DMM. Comparing continuous and perfusion cultivation of microalgae on recirculating aquaculture system effluent water. Bioresource Technology. 2025;418:131881.

26. Hawrot-Paw M, Koniuszy A, Gałczyńska M, Zając G, Szyszlak-Bargłowicz J. Production of microalgal biomass using aquaculture wastewater as growth medium. Water. 2020;12(1):106.

27. Borg-Stoveland S, Draganovic V, Spilling K, Gabrielsen TM. Successful growth of coastal marine microalgae in wastewater from a salmon recirculating aquaculture system. Journal of Applied Phycology. 2024.

28. Santin A, Russo MT, Ferrante MI, Balzano S, Orefice I, Sardo A. Highly valuable polyunsaturated fatty acids from microalgae: strategies to improve their yields and their potential exploitation in aquaculture. Molecules. 2021;26(24):7697.

29. Shahar B, Haim E, Kuc ME, Azerrad SP, Dudai N, Kurzbaum E. Simplified and cost-effective modulatory photobioreactor setup for upscaling microalgal culture for research and semi-industrial purposes. Algal Research. 2023;74:103200.

30. Guillard RR. Culture of phytoplankton for feeding marine invertebrates. Culture of marine invertebrate animals: Springer; 1975. p. 29–60.

31. Kérouel R, Aminot A. Fluorometric determination of ammonia in sea and estuarine waters by direct segmented flow analysis. Marine Chemistry. 1997;57(3– 4):265–75.

32. Spilling K, Seppälä J. Measurement of Fluorescence for Monitoring Algal Growth and Health. In: Spilling K, editor. Biofuels from Algae: Methods and Protocols. New York, NY: Springer New York; 2020. p. 41–5.

33. Team P. RStudio: Integrated Development Environment for R. Posit Software, PBC, Boston, MA. 2023.

34. Chuka-ogwude D, Ogbonna J, Borowitzka MA, Moheimani NR. Screening, acclimation and ammonia tolerance of microalgae grown in food waste digestate. Journal of Applied Phycology. 2020;32:3775–85.

35. Collos Y, Harrison PJ. Acclimation and toxicity of high ammonium concentrations to unicellular algae. Marine pollution bulletin. 2014;80(1-2):8–23.

36. Zhao P, Gu W, Huang A, Wu S, Liu C, Huan L, et al. Effect of iron on the growth of Phaeodactylum tricornutum via photosynthesis. Journal of Phycology. 2018;54(1):34–43.

37. Danouche M, El Ghatchouli N, Arroussi H. Overview of the management of heavy metals toxicity by microalgae. Journal of Applied Phycology. 2022;34(1):475– 88.

38. Zhou B, Ma J, Chen F, Zou Y, Wei Y, Zhong H, et al. Mechanisms underlying silicon-dependent metal tolerance in the marine diatom Phaeodactylum tricornutum. Environmental Pollution. 2020;262:114331.

39. Wang C, Li J, Li S, Lin S. Effects and mechanisms of glyphosate as phosphorus nutrient on element stoichiometry and metabolism in the diatom Phaeodactylum tricornutum. Applied and Environmental Microbiology. 2024;90(2):e02131–23.

40. Zhao P, Gu W, Wu S, Huang A, He L, Xie X, et al. Silicon enhances the growth of Phaeodactylum tricornutum Bohlin under green light and low temperature. Scientific Reports. 2014;4(1):3958.

41. Yi Z, Su Y, Cherek P, Nelson DR, Lin J, Rolfsson O, et al. Combined artificial high-silicate medium and LED illumination promote carotenoid accumulation in the marine diatom Phaeodactylum tricornutum. Microbial Cell Factories. 2019;18(1):209.

42. Barta DG, Coman V, Vodnar DC. Microalgae as sources of omega-3 polyunsaturated fatty acids: Biotechnological aspects. Algal Research. 2021;58:102410.

43. Vázquez-Romero B, Perales JA, de Vree JH, Böpple H, Steinrücken P, Barbosa MJ, et al. Techno-economic analysis of microalgae production for aquafeed in Norway. Algal Research. 2022;64:102679.

44. Santin A, Balzano S, Russo MT, Palma Esposito F, Ferrante MI, Blasio M, et al. Microalgae-based PUFAs for food and feed: current applications, future possibilities, and constraints. Journal of Marine Science and Engineering. 2022;10(7):844.

45. Tripathi R, Gupta A, Thakur IS. An integrated approach for phycoremediation of wastewater and sustainable biodiesel production by green microalgae, Scenedesmus sp. ISTGA1. Renewable Energy. 2019;135:617–25.

46. Malibari R, Sayegh F, Elazzazy AM, Baeshen MN, Dourou M, Aggelis G. Reuse of shrimp farm wastewater as growth medium for marine microalgae isolated from Red Sea–Jeddah. Journal of Cleaner Production. 2018;198:160–9.

